# Whole genome comparisons reveal panmixia among fall armyworm (*Spodoptera frugiperda*) from diverse locations

**DOI:** 10.1101/2020.09.25.314005

**Authors:** Katrina A. Schlum, Kurt Lamour, Caroline Placidi de Bortoli, Rahul Banerjee, Scott J. Emrich, Robert Meagher, Eliseu Pereira, Maria Gabriela Murua, Gregory A. Sword, Ashley E. Tessnow, Diego Viteri Dillon, Angela M. Linares Ramirez, Komivi S. Akutse, Rebecca Schmidt-Jeffris, Fangneng Huang, Dominic Reisig, Juan Luis Jurat-Fuentes

## Abstract

The fall armyworm (*Spodoptera frugiperda* (J.E. Smith)) is a highly polyphagous agricultural pest with long-distance migratory behavior threatening food security worldwide. This pest has a host range of >80 plant species, but two host strains are recognized based on their association with corn (C-strain) or rice and smaller grasses (R-strain). In this study, the population structure and genetic diversity in 55 *S. frugiperda* samples from Argentina, Brazil, Kenya, Puerto Rico and the United States (USA) were surveyed to further our understanding of whole genome nuclear diversity. Comparisons at the genomic level suggest panmixia in this population, other than a minor reduction in gene flow between the two overwintering populations in the continental USA that also corresponded to genetically distinct host strains. Two maternal lines were detected from analysis of mitochondrial genomes. We found members from the Eastern Hemisphere interspersed within both continental USA overwintering subpopulations, suggesting multiple individuals were likely introduced to Africa. Comparisons between laboratory-reared and field collected *S. frugiperda* support similar genomic diversity, validating the experimental use of laboratory strains. Our research is the largest diverse collection of United States *S. frugiperda* whole genome sequences characterized to date, covering eight continental states and a USA territory (Puerto Rico). The genomic resources presented provide foundational information to understand gene flow at the whole genome level among *S. frugiperda* populations.

## Introduction

Larvae of the fall armyworm, *Spodoptera frugiperda* (J.E. Smith, 1797) (Lepidoptera: Noctuidae), are highly polyphagous agricultural pest affecting key food and fiber staples such as corn (*Zea mays* L.), cotton (*Gossypium spp.* L.), sorghum (*Sorghum bicolor* L.), rice (*Oryza sativa* L.), and vegetable crops [1]. Damage by *S. frugiperda* in its native subtropical range in the Americas resulted in 15-100% yield loss, depending on the level of infestation [2]. In most of the continental USA this pest does not diapause or survive cold winters. Overwintering populations in southern Texas and Florida migrate yearly northward over several generations to populate northern regions in USA and Canada [3].

For more than a decade, effective control of *S. frugiperda* in the Western hemisphere has been provided mainly by genetically modified corn and cotton producing insecticidal Cry and Vip3A proteins from the bacterium *Bacillus thuringiensis* (*Bt*). However, resistance to Cry1F, Cry1Ab and Cry1A.105 insecticidal proteins quickly developed in Puerto Rico and the continental USA (Florida and North Carolina) [4, 5], as well as Brazil [6] and Argentina [7]. More recently, the economic importance of this pest has further increased with its introduction in sub-Saharan Africa [8], subsequent spread to India and Southeastern Asia [9], and more recently Australia [10]. An estimate using data from twelve African countries indicates that yield losses resulting from *S. frugiperda* injury could be 21-53% of their annual corn production, which equals to a loss of US$2.5-$6.2 billion in losses [11]. While yet to be detected in molecular screens [12], resistance alleles to *Bt* toxins could have been carried by invasive *S. frugiperda* to the Eastern hemisphere.

Populations of *S. frugiperda* are composed of sympatric mixtures of two genetically differentiated strains based on host preference, a “rice” (R) strain feeding on rice, millet and smaller grasses, and a “corn” (C) strain feeding preferentially on corn and sorghum [13, 14]. There is evidence supporting that this differentiation involves reproductive incompatibility [15] and differential susceptibility to xenobiotics [16]. Signatures of positive selection for genes involved in chemoreception, detoxification, and digestion were also detected in whole-genome comparisons of nine C- and R-strain individuals [17]. Being morphologically indistinguishable, C and R individuals are discriminated using genetic markers located on the mitochondrial *COI* (cytochrome oxidase subunit I) and sex-linked (Z chromosome) *Tpi* (*triosephosphate isomerase*) genes [18-21]. However, host preference is not absolute and discrepancies among haplotype markers have been reported. For example, the *Tpi* marker agrees with assortative mating and host assessments [22] in describing *S. frugiperda* samples from corn in the Eastern Hemisphere as C-strain, yet a predominant *COI*-R strain marker is obtained in these collections [9]. These conflicting results may reflect interstrain hybridization [22] or be driven by maternally inherited symbionts that skew the distribution and diversity of certain haplotypes [23]. Additional factors influencing these conflicting haplotyping results may include incomplete lineage sorting and plant host behavioral plasticity.

Nucleotides found at *COI* host strain marker sites have been commonly grouped into “haplotypes” with relative proportions associated to geographic origin [24], enabling differentiation of “Texas” and “Florida” overwintering *S. frugiperda* populations in the USA. Migratory studies with these haplotypes determined that the “Texas” population is distributed throughout South, Central and North America [25, 26], while the “Florida” population locates to Florida and the Caribbean but migrates through the eastern USA seaboard to reach Canada [27, 28]. Haplotype profiling of *S. frugiperda* from Africa, India and Southeastern Asia supports introductions in Africa of individuals from the “Florida” population and subsequent spread to Asia [9, 12].

In this work, we collected and sequenced 55 genomic DNA (gDNA) samples of *S. frugiperda* from three continents, with an emphasis on C-strain individuals from North and South America. Genome-wide comparisons allowed testing for gene flow between geographically distant *S. frugiperda* populations and comparing genetic diversity between field and laboratory-reared *S. frugiperda.* We find that based on F_st_ values there is low genetic differentiation between Texas vs. Florida and that samples from these locations display similar genetic nucleotide diversity. Host strain differentiation was supported by nuclear and mitochondrial genome differences. Comparison of mitochondrial genomes detected two clusters where the clusters correlate by and large with host strain and resistance to the Cry1F toxin. The exceptions to this correlation suggest that C-strain individuals based on nuclear genome have likely mated with R-strain individuals. We also observed no detectable reduction in genetic diversity in well-established *S. frugiperda* laboratory-reared colonies, which supports using laboratory strains of this pest as a model for field populations.

## Methods

### Samples and strain typing

Details of the 55 laboratory and field-collected *S. frugiperda* samples analyzed are found in Supplementary Table 1. Adults (moths) were captured using sex pheromone baited traps [29], with most collection sites near corn plantings to optimize trap capture efficiency of C-strain males. Laboratory-reared samples were obtained from rearing facilities. Susceptibility to Cry1F toxin from *B. thuringiensis* for these laboratory samples was tested in bioassays presented elsewhere [5, 30-32]. Larval samples were collected at field and laboratory rearing locations and used when available. The collected specimens were identified as *S. frugiperda* by morphology features [33] and stored at -20°C until required for analysis.

We followed a sample naming protocol that included the first three letters representing the country of origin (Bra for Brazil, Arg for Argentina, USA for United States of America, Ken for Kenya, and Pue for Puerto Rico), the next two letters representing the first two letters of the state/province of origin (SP for Sao Paulo, TX for Texas, etc. XX when unknown), and the third letter representing the Cry1Fa susceptibility phenotype when known (“r” for resistant, “s” for susceptible and “u” for unknown). A number was used when necessary to differentiate samples with the same geographic origin and Cry1F susceptibility phenotype.

Host strain (C versus R) was determined based on sequence identity at specific marker positions in reference mt*CoI*1164 and *Tpi*183 sequences, as described elsewhere [12]. The mitochondrial *COI1287* marker identified all samples as C-strain, so the nuclear *Tpi*183 marker was used as a more reliable marker of host strain [34]. The mitochondrial *COI*1164 marker was not used because it did not fall under any of the pre-defined *COI* haplotypes [34] for any of the samples.

### DNA extraction, library preparation, and sequencing

Genomic DNA was isolated from individual carcasses of fifth instar *S. frugiperda* larvae after dissecting the gut tissue or from legs or heads of adults using the Pure Link Genomic DNA mini kit (Invitrogen), following manufacturer’s protocols, and then quantified using a Nanodrop spectrophotometer (Thermo Scientific). Extracted DNA was then sheared randomly to between 250-500 bp using a Covaris M220 focused ultrasonicator (Woburn, MA), according to manufacturer’s instructions. The fragmented DNA was then ligated with dual-indices using a KAPA Hyper prep PCR-free library kit (Roche) according to the manufacturer’s directions. The ligated fragments were quantified using quantitative PCR and a KAPA Library quantification kit (Roche), and then submitted for sequencing on an Illumina HiSeqX device running a 2 x 150 bp paired-end configuration (Admera Health, NJ). Raw paired-end sequence reads for each sample (55) are available for download in the NCBI Sequence Read Archive (SRA) under SRR12044614-SRR12044668 with associated metadata available under NCBI BioProject id PRJNA640063. The raw data was processed to remove low quality reads using the CLC Genomics Workbench v9.5.2 (Qiagen) trim function with default parameters.

### Filtering and mapping

Raw reads were quality-trimmed at both ends, filtered for adapter sequences and error corrected for known Illumina artifacts and PhiX sequencing control using BBDuk (https://jgi.doe.gov/data-and-tools/bbtools/) with options trimq = 15 and filter = 23. Further quality control and confirmation that adapter and non-relevant (contaminants, primer artifacts) sequences were removed from filtered reads were performed using FastQC [35]. Given the prevalence of C-strain among our samples and similar number of variants called when using both *S. frugiperda* corn and rice genomes [36], we mapped the remaining reads to the *S. frugiperda* corn reference genome (v3.1) downloaded from https://bipaa.genouest.org/sp/spodoptera_frugiperda_pub/ (assembled length of 312 Mb across

29,949 scaffolds [17]) using a Burrows-Wheeler aligner algorithm (bwa-mem) [37]. Variants were then called using the SAMtools mpileup utility [38], resulting in detection of 126,977,977 single nucleotide polymorphisms (SNPs) and indels. The unfiltered dataset contained 120,398,863 SNPs. Three samples (USATXu1, BraSPr2 and BraSPr1) were removed from SNP analysis because they had missing variant call rate greater than 50%, resulting in 52 samples being used for SNP PCA-based analyses. For nucleotide diversity analyses, SNPs were filtered using VCFtools [39] based on both minor allele frequency being less than or equal to 0.05 and presence in at least 50% of the 52 samples surveyed. This filtering further reduced SNP number to 204,784.

### Phylogenomics

The Mash distance method was used to measure pairwise dissimilarity of genomic sequences [40]. One key advantage of sketching-based approaches is that they neither require *de novo* genome assembly nor a pre-existing reference genome to identify related individuals. The Mash method estimates sequence similarity via the Jaccard similarity coefficient (the ratio of shared k-mers to total k-mers) for a “random” subset of k-mer pairs for each genome surveyed. The Jaccard similarity (J) is then used to calculate dissimilarity between two genomes (*D*) as D = 1-J. All *S. frugiperda* individuals were sketched with Mash [40] using a 10,000 sketch size, k-mer size of 21 and minimum copy of each k-mer equal to 2, per previous recommendations of representing genomes with minimal computational costs [40]. All sample sketches were screened against the Mash Refseq database and other *Spodoptera* species (*Spodoptera exigua*, GCA_011316535.1; and *Spodoptera litura*, GCA_002706865.2) as outgroups.

### SNP diversity

Weighted F_st_ values as indicators of total genetic variance in a subpopulation were calculated using VCFtools [39] by the Weir and Cockerham estimator on 1 kbp, 3 kbp and 5 kbp windows. The variation in window size was used to confirm that there was little to no change in F_st_ when the window was varied. For F_st_ calculations using the mitochondrial genome, each base pair position was compared. Nuclear genome (pi) values were calculated using 1 kbp windows when comparing corn vs. rice host strain, and 3 kbp when comparing F_st_ values for overwintering FL/TX vs. USA laboratory samples. We also used 3 kbp windows for F_st_ comparisons across the nuclear genome with populations defined by mitochondria haplotype network groupings, Mash genome defined “haplotypes”, and host strain haplotypes. For all nucleotide diversity analyses, a total of 3,558,854 SNPs were used to assess diversity using VCFtools [39].

### Mitochondrial genome assembly

Mitochondrial sequences were extracted from the whole genome sequence reads using NOVOplasty [41] with the mitochondrial partial *COI* sequence MH932092.1 as the seed. A continuous complete mitochondrial chromosome of around 15 kbp was generated for all 55 samples, except for BraBAr5, which was excluded from further analysis. The whole mitochondrial FASTA sequences were assembled for each sample and then aligned via bwamem (https://github.com/lh3/bwa) with default options. Mitochondrial variants were called using SAMtools mpileup based on the *S. frugiperda* corn reference genome at LepidoDB (https://bipaa.genouest.org/sp/spodoptera_frugiperda_pub/). The vcf files were compressed using SAMtools bgzip, indexed using SAMtools tabix and merged using SAMtools BCFtools merge [38]. Missing genotypes were replaced with sequence at the mitochondrial reference genome from the *S. frugiperda* corn genome at LepidoDB. The vcf files were then filtered by removing 16 multi-allelic sites and 92 uninformative sites due to being found in 2 samples or less via the informloci function in the adegenet R package [42], resulting in 298 sites remaining. These remaining mitochondrial sites were used to build a haplotype network using the R package pegas haploNet function [43] with Euclidean distance and an infinite site model used for building the network.

### Structure

PLINK –make-bed [44] was used to generate the input file to FastStructure [45], a variation of Structure made for larger SNP datasets. FastStructure is a generative model-based approach based on Hardy-Weinberg equilibrium assumptions between alleles and linkage disequilibrium between genotyped loci. The FastStructure script structure.py was used to determine k, the number of assumed populations or genetic groups that share a subset of allele frequencies from k = 2 to 10. Then k =2 was chosen by the FastStructure’s choosek.py script for model complexity that maximizes marginal likelihood, and k = 1 for model components to explain structure. Additionally, PCA eigenvalues and eigenvectors were generated using PLINK 2.0 [44, 46] and visualized in R using ggplot2 [47].

## Results

### Host strain typing

The majority (26, 47.3%) of the 55 *S. frugiperda* samples used in this study originated from field collections representing eight states in the USA. The average mapping rate for all samples was 88.2% with a range of between 60.62% to 96.82% range per SAMtools flagstat [38]. Identification of the C- and R-host strains for each individual was performed using nuclear *Tpi* and mitochondrial *COI* genetic markers described elsewhere [26]. Given that sample collections were from or around cornfields, we expected to observe a majority of C-strain samples. The *COI*1164 marker identified all samples as C-strain. In contrast, typing based on the nucleotides at the *TPi*183 site resulted in 42 samples identified as C-strain individuals (C/C), 8 as R-strain individuals (T/T), 3 as interstrain hybrids (C/T) and 2 that were undetermined due to low sequencing coverage at this locus. All eight identified R-strain samples were from the USA (3 from Texas, 2 from Maryland and Tennessee each, and 1 from Puerto Rico).

### Whole genome diversity, nucleotide diversity and differentiation

The FastStructure tool was used first to determine possible population structure among the samples. The number of groups (k) was analyzed for k = 1 to k = 10, but all samples clustered as one group even when the number of groups was increased. This lack of clear population structure was further confirmed using principal component analysis (PCA) on a filtered set of 204,784 nuclear SNPs (see Methods). PC1 through PC10 was plotted and no structure was found based on country of origin, host strain or susceptibility to the Cry1F insecticidal protein from *Bt* (data not shown for PC2 to PC10). We noted that PCA using the first two principal components uncovered weak substructure within the USA and Brazil samples, where the majority of samples clustered in a larger cluster and four samples (USAFLu9 from Florida, and BraMGr2, BraBAr2, and BraMGr5 from Minas Gerais in Brazil) grouped together in a smaller cluster with 1 Florida sample (USAFLr2) the most differentiable from the two clusters (Fig. 1). All these small cluster samples except USAFLu9 are laboratory samples, and therefore this grouping may be a result of the side effect of genetic drift associated with the colony formation process.

**Figure 1:**
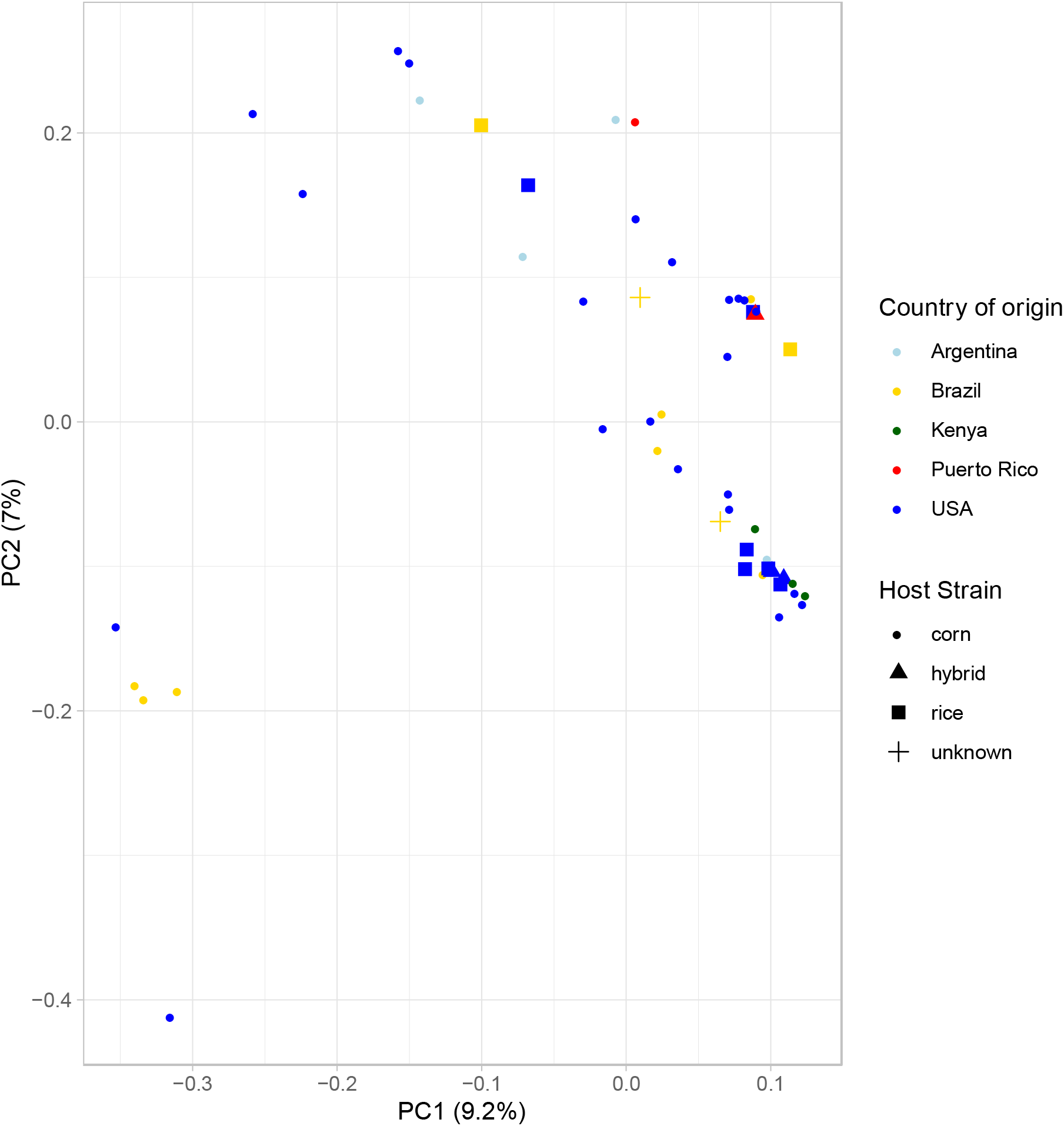
Principal component analysis (PCA) plot of 52 *Spodoptera frugiperda* samples (3 samples removed due to missing rate ≥ 50%) based on 3,558,854 SNPs. Country of origin and host strain as determined by the nuclear *Tpi* marker are visualized by color and shape, respectively, as shown in the legend.

Given the overall lack of clear population structure observed based on origin, we next applied genome sketching [40] and traditional distance-based hierarchical clustering as a third strategy to analyze these samples. The hierarchical cluster dendrogram defined by the genome-wide distance estimates based on the fraction of shared k-mers between each pair of genomes was cut at height 0.05 as it resulted in at least two primary clusters (Texas vs. non-Texas) among the samples analyzed (Fig. 2). This cut height generated 8 clusters, 6 containing less than 3 samples and were merged to the major cluster, while the remaining 10 samples were left in the remaining 2 minor cluster. The first cluster was comprised of samples from the USA (USATXu2-5 from Texas, USATNu1-2 from Tennessee, USAMDu1-2 from Maryland, USAFlu2 from Florida and PueSlu1 from Puerto Rico), while the second cluster included individuals from all countries sampled with the majority of samples from USA representing Florida/eastern seaboard. The three samples from Puerto Rico were interspersed across both clusters and always clustered with Florida samples, yet genomic distances between clusters were small (< 0.05 Mash distance). Similar to the PCA clusters, these two clusters did not correlate with host strain designation or geographic origin, with the first cluster including both R- and C-strain samples and the second cluster containing both R- and C-strain samples as well as three interstrain hybrids (Supplemental Fig. S2).

**Figure 2:**
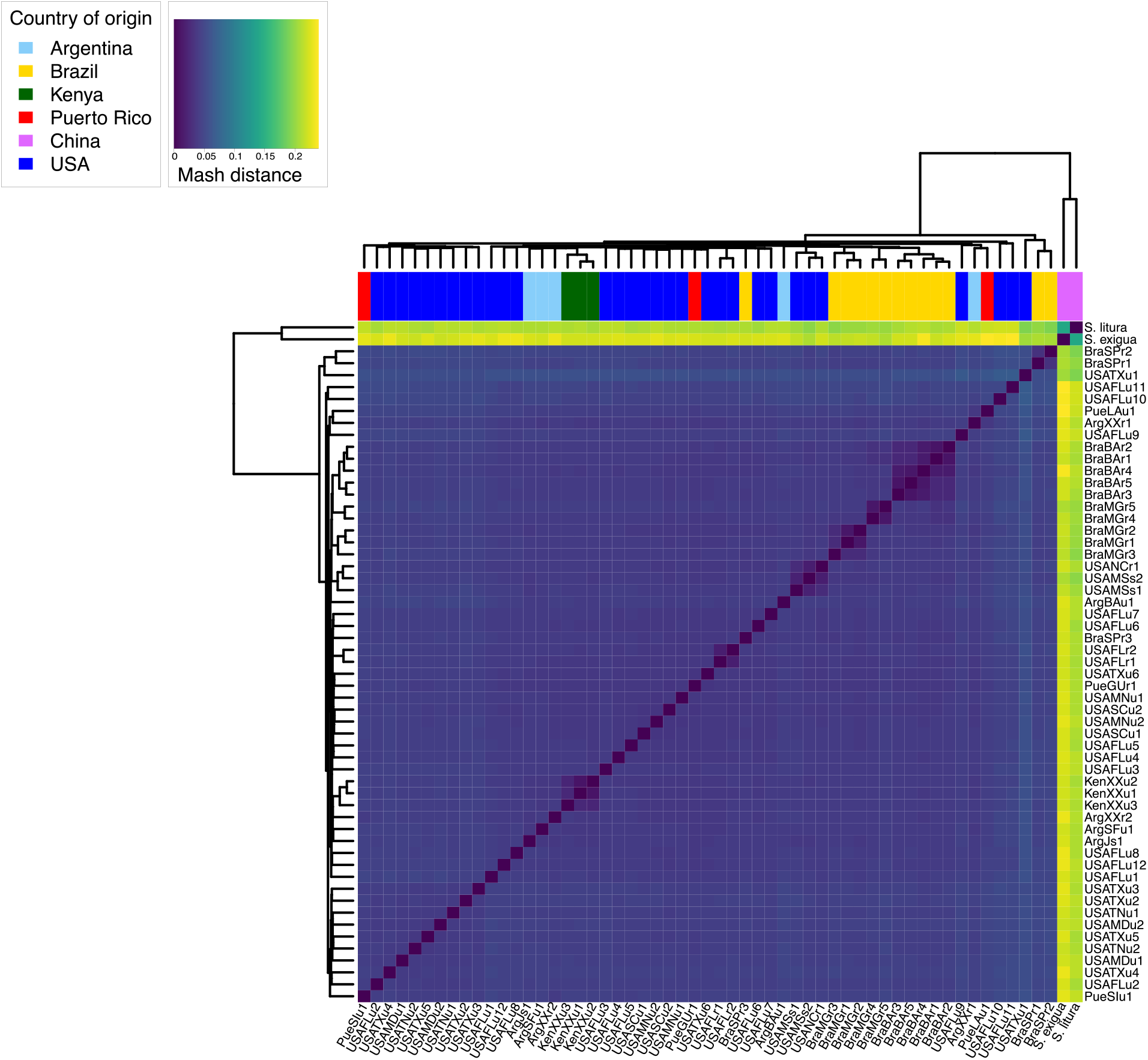
Dissimilarity heatmap of Mash distances across all 55 *Spodoptera frugiperda* samples, colored by country of origin as indicated. Two outgroups (*Spodoptera litura* and *Spodoptera exigua*) were added. Samples were named as described in Materials and Methods.

Mash distance is a proxy for genome pairwise distance, with 0 representing identical genomes (all k-mers that comprise the sketches are present in both sequences). The Mash distances across all countries surveyed were averaged to be <0.05 (Table S2, range of average distance was 0.043 to 0.045), probably reflecting recent population expansion. Moreover, these Mash distances support that R and C-strains are members of the same species, as the estimated genetic variation is comparable to at least 95% average nucleotide identity (ANI), which is the ANI cutoff for eukaryotic species [40].

### Gene flow between overwintering populations in the USA

Previous work based on the ratios of four pre-determined *COI* haplotypes suggested that the overwintering Florida *S. frugiperda* population is mostly reproductively isolated from the overwintering population in Texas, with genetic exchange mostly occurring at the north and south ends of the Appalachian mountain chains [27]. As mentioned above, we observed that C-strain samples from Texas and Florida/eastern seaboard did not separate based on hierarchical clustering of genome-wide distances, and Florida samples were found across both groups (Fig. 2). Further, when comparing the two R-strain samples from Texas and the fourteen C-strain samples from Florida/eastern seaboard, we detected a low pairwise weighted F_st_ value (0.0111) based on 3,558,854 nuclear sites. This very low F_st_ value suggests no population separation at the nuclear level, supporting recent population separation or ongoing gene flow between Texas and Florida overwintering populations, independently of host strain.

Based on the same Texas R-strain and Florida C-strain groupings, mean diversity was determined. Mean diversity for the field samples from Texas R-strain and Florida C-strain samples was estimated at 0.0631 and 0.0556, respectively (Fig. 3). Next, we determined the F_st_ value based on 406 mitochondrial SNPs found across all samples in these groups. The mitochondrial F_st_ for Texas R-strain and Florida C-strain samples was estimated at 0.7961. This F_st_ estimate being close to 1 suggests genetic differentiation between the R and C strains is the strongest at the mitochondrial level, possibly due to two main ancestral lineages. Although we see a stronger difference between geographic regions than between R and C strains in this study, the estimated mitochondrial to nuclear F_st_ ratio of 46.8 is consistent with previously reported sex-based differences in dispersal [48] and/or mating preference differences between strains [49]. Even so, our data suggest hybridization occurs and some C-strain nuclear genomes have a clearly R-strain associated mitochondrial genome (see Fig. 5).

**Figure 3:**
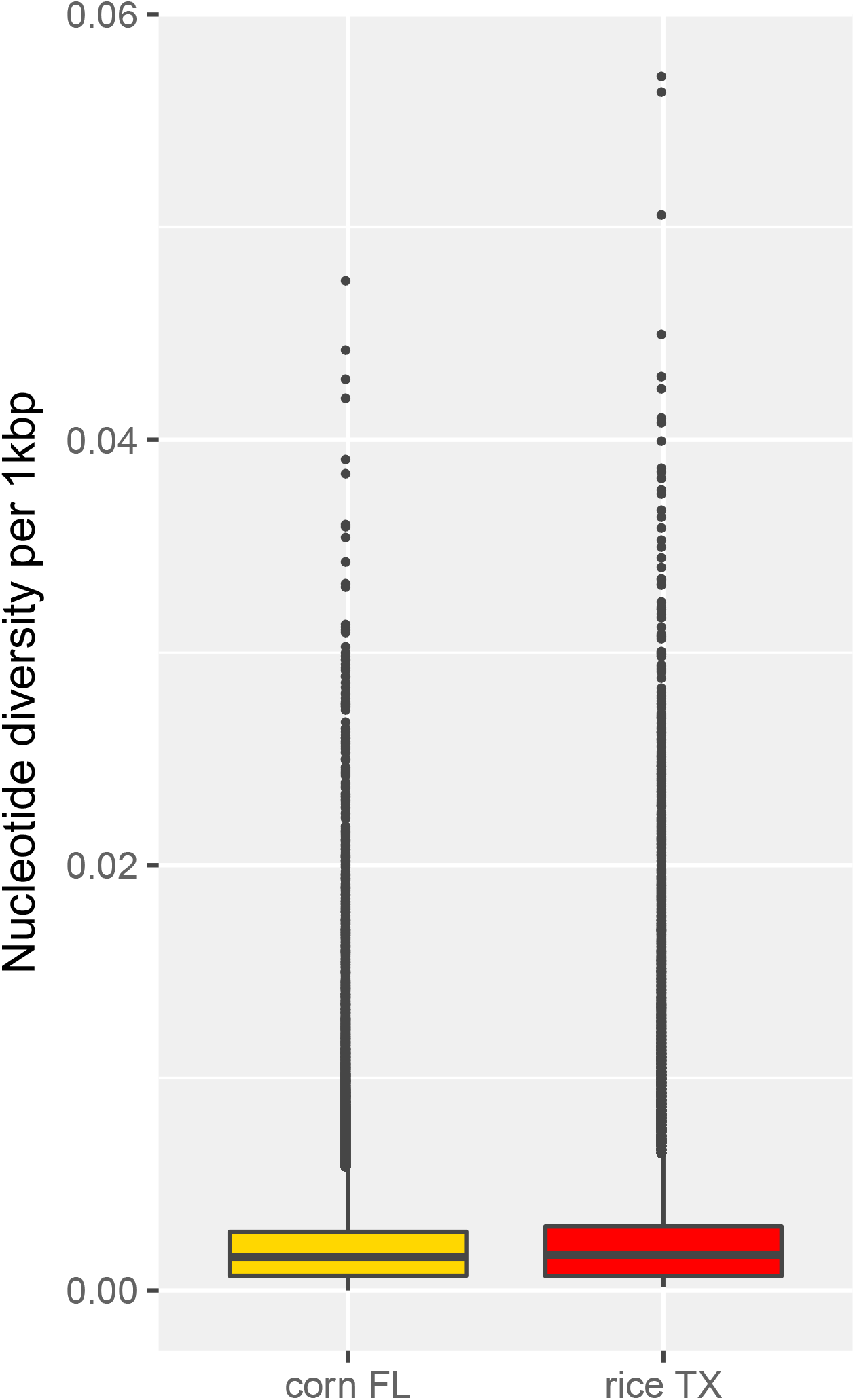
Nucleotide diversity box plots between C-strain Florida (FL) vs. R-strain Texas (TX) samples.

**Figure 4:**
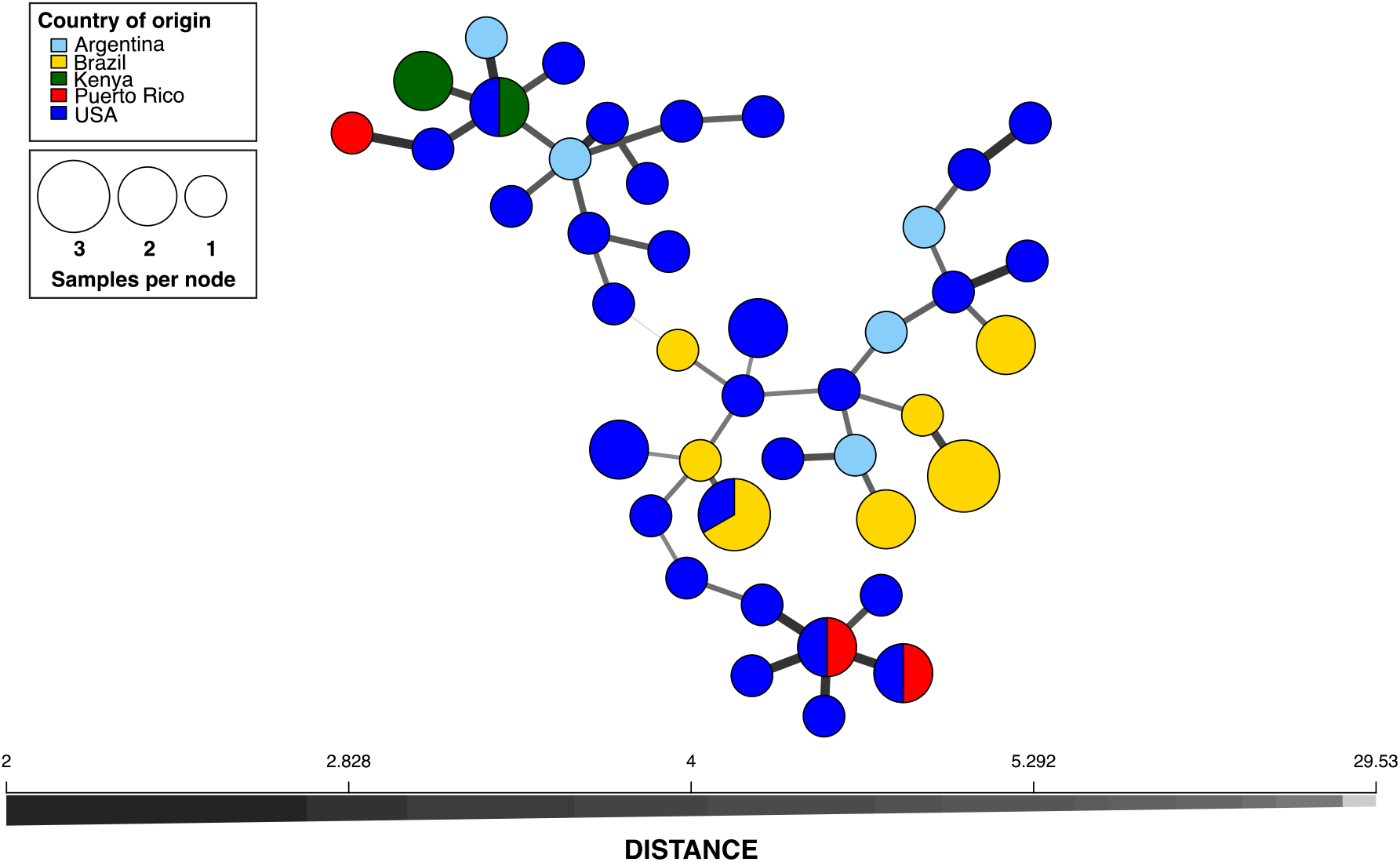
Mitochondrial haplotype network of all 54 *S. frugiperda* mitochondrial chromosome level assembled individuals, colored by country of origin.

**Figure 5:**
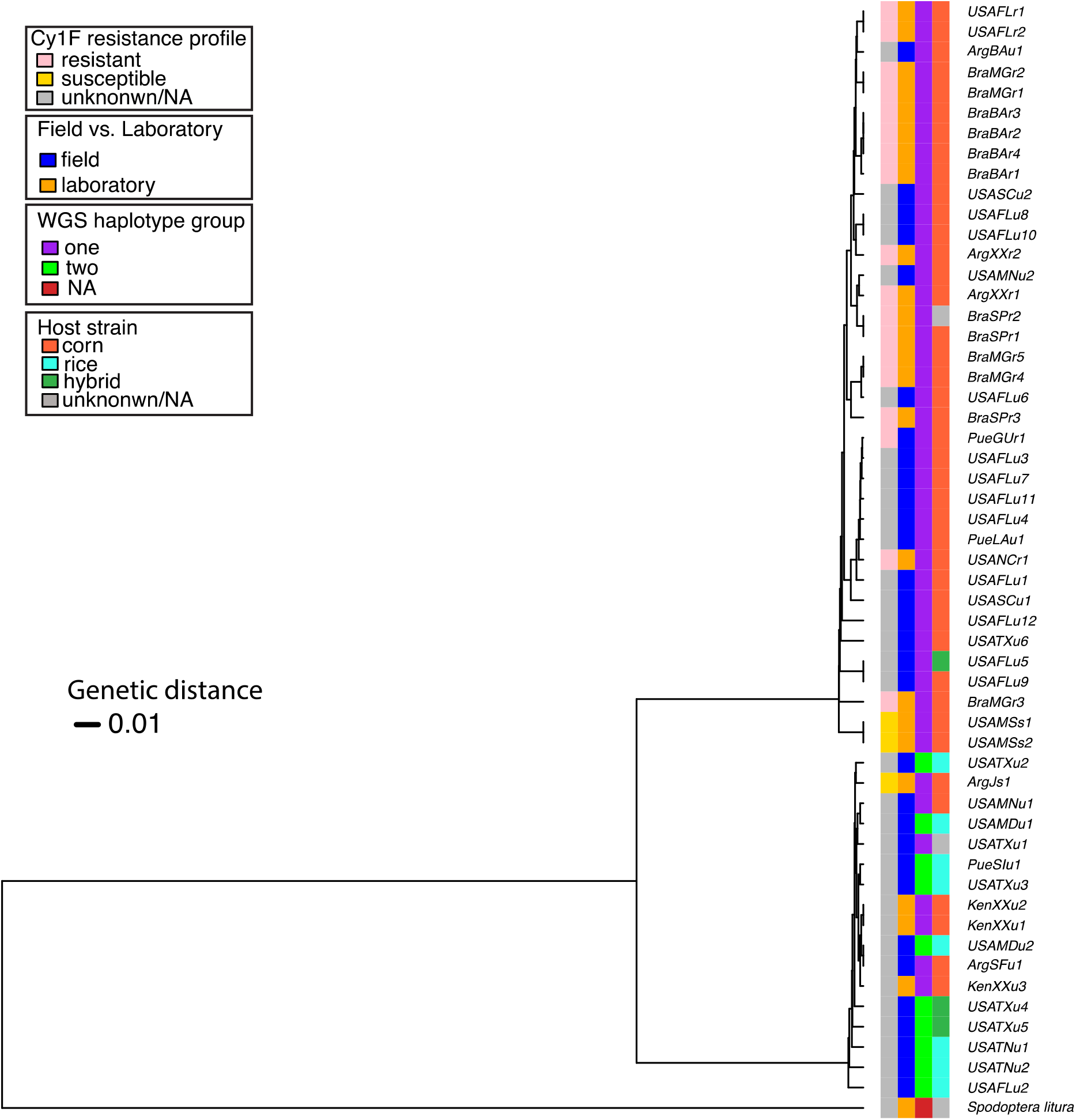
UPGMA distance-based tree on biallelic variant SNPs in 54 mitochondrial chromosome level assembled individuals rooted with *Spodoptera litura* as an outgroup.

### Laboratory vs. field diversity

Laboratory-reared strains of *S. frugiperda* are commonly used for diverse physiological, toxicological and genetic studies, yet their genetic resemblance to field populations has not been investigated. To investigate the differentiation level between two well established laboratory strains (>20 years) originating from the same locations in the USA (Mississippi, MS) and all field USA populations surveyed, we estimated a weighted F_st_ (fixation index). As shown in Table 1, there was moderate differentiation between the MS laboratory strains and field USA field populations, with an F_st_ range of 0.0056 to 0.1109. Moderate differentiation was also observed when comparing the MS laboratory strains with other more recently established (<10 years) laboratory populations from North Carolina (NC) and Florida (FL), with an F_st_ range of 0.0024 to 0.0874. The samples least differentiated (F_st_ = 0.0631) from the MS laboratory populations were collected from TX/MN locations, while the most distinct (F_st_ = 0.1263) were the samples from Maryland (MD). Overall, these moderate F_st_ values [50] suggest small differences at the allelic level between laboratory and field-collected groups. Genetic diversity in the two laboratory MS colonies was slightly different but comparable to diversity in field USA populations (F_st_ = 0.0981 on average) and to either of the two overwintering populations (F_st_ = 0.0594).

Additionally, we surveyed overall genomic diversity using 3 kbp windows and estimated mean diversity for the same two MS laboratory strains compared to all field and laboratory *S. frugiperda* samples surveyed. Mean nuclear diversity was 0.0011 for Benzon (USAMSs1) and SIMRU (USAMSs2) laboratory strains (n=2), compared to 0.0015 for field Texas samples (n=7), and 0.0015 for field Florida/eastern USA samples (n=14).

### Diversity across the mitochondrial genome

We investigated the maternal lineage among all but one of the *S. frugiperda* samples (due to lack of a complete mitochondrial genome in sample BraBAr4) by comparing genetic diversity of whole mitochondrial (mt) genomes. Using a total of 390 SNPs, we generated a haplotype network as described in Materials and Methods (Fig. 4). Consistent with a large amount of standing variation in the species, even the mitochondrial sequences appeared unique with all but eight samples forming a unique haplotype. These exceptions included three haplogroups of only Brazil samples, a second group of Kenya and USA samples, and a third haplogroup of samples from Kenya and two haplogroups of Florida, USA and Puerto Rico samples (Fig. 4). There were two main “haplotype” clusters that separated in the network, with average genetic difference between the two clusters of 29.53, supporting evidence for two main maternal lineages within the samples surveyed. Using these two haplotype clusters as labels, we mapped the mt haplotype clusters to the two whole genome sequence (WGS) “haplotype” clusters found in Mash distance-based tree against a mitochondrial variant UPGMA distance-based tree (Fig. 5). In accordance with the previously estimated F_st_, the mt and WGS clusters did not correlate completely. However, all samples in one of the mt haplotypes clustered in one of the WGS haplotypes (“one” in purple in Fig. 5), while the other mt haplotype contained a mix of WGS haplotypes. Interestingly, and while not all samples were phenotyped, all samples classified as resistant to the Cry1F toxin clustered in one of the mt haplotype groups.

### Discussion

We report on the genome-level comparison of 55 *S. frugiperda* samples from diverse locations, mostly from the native range in the Americas but also including samples from an African location (Kenya) where the insect is a devastating exotic pest. While using the C-strain *S. frugiperda* genome for alignments could have underestimated diversity in R-strain individuals, this approach is supported by previous surveys using both corn and rice genomes with minimal differences found based on the number of variants called and F_st_ values [36].

The population structure of the 55 genomic *S. frugiperda* samples was surveyed using parametric and non-parametric methods. Based on FastStructure (parametric), no population structure was found either within samples from the Eastern Hemisphere or between the Eastern and Western Hemisphere samples. Since methods based on parametric estimation of allele frequencies methods are sensitive to sample size, one possibility is that the lack of structure detected is a reflection of the relatively small number of R-strain samples (only 8 individuals and potentially 2 hybrids) analyzed. A more even distribution would address potential limited sampling of some locations surveyed, especially samples representing the Eastern Hemisphere (e.g. Africa).

Although PCA based on the nuclear genome agreed with the little to no population structure as found by FastStructure, the Mash distance-based analysis on the nuclear genome and SNP Hamming-based distance method based on the mitochondrial genome detected two subpopulations within the samples surveyed, which associate with the previously speculated two subpopulations in the continental U.S. based on four mitochondrial *COI* haplotypes [51]. The two clusters clearly indicate that the corn and rice strain individuals are more similar locally than between geographic areas, suggesting some level of hybridization between host strains in the USA. Further, we have analyzed individuals that appear to have a C-strain nuclear genome with an R-strain associated mitochondria (maternal lineage). This is consistent with the conflicting host strain assignments using specific genetic markers between R-strain and interstrain hybrids that was previously reported [22]. We still observe host differentiation at the nuclear genome level, though, and may suggest current single locus markers are not sufficient for surveying all *S. frugiperda*.

Our analysis included samples collected from locations (Maryland & Tennessee) close to predicted hybridization zones for TX and FL overwintering *S. frugiperda* populations [27], and as expected these associated with both geographic regions. There were also three exceptions to the geographic separation: two Texas samples were found in the predominately Florida represented group and one Florida sample was found in the predominately Texas population. This observation suggests that the reproductive isolation between Texas and Florida/Eastern seaboard is not absolute and mating may be occurring in overlapping locations, as previously suggested [27]. Future work should include increased sampling from potential hybrid zones (Georgia/New York) and Texas. The Mash distance-based clusters suggested that Puerto Rico samples are closer to Florida, since they were found in sub-clusters with Florida-based samples. However, due to low representation and small genomic distances between each cluster, our data is not able to show conclusive support for migration models suggesting significant exchanges between *S. frugiperda* from Puerto Rico and Florida [28].

The low amount of genetic variation across the whole genome between the C- and R-strains is in line with previous reports, [17] and suggest that the genome differentiated after the ecological differentiation of host strains or that barriers are isolating C- and R-strains (i.e. prezygotic reproductive isolation or mating preference). In testing this hypothesis, future research should involve analyses of a larger sample size of C- and R-strain individuals from Texas and Florida.

Mitochondrial F_st_ estimates of 0.7961 obtained between C- and R-host strains were highly similar to the F_st_ value (0.938) from a prior analysis of 9 corn and 9 rice sympatric samples from Mississippi (USA) [17], and the nuclear DNA F_st_ found in our study (0.0173) is also similar to the previous study’s estimate (0.019). The two mitochondrial haplotypes found in our analysis could be explained by selective mating or some form of reproductive incompatibility. Although susceptibility to Cry1F was not known for all samples, we observed that all Cry1F-resistant strains clustered to one of the mitochondrial haplotypes, despite their geographic origin. While speculative given the comparatively low number of confirmed Cry1F-resistant samples, this grouping may reflect partitioning of resistance with the haplotype group only containing C-strain samples. Further work including Cry1F-resistant R-strain samples would be necessary to test for associations between mitochondrial haplotype and resistance to Cry1F.

One possibility to explain the mitochondrial haplotypes detected could be that they are based on reproductive incompatibility from infection by parasites such as *Wolbachia* [52]. However, we only detected short fragments (<100 bp) matching to the RefSeq *Wolbachia* genomes on NCBI as of May 9, 2020 using BLAST (data not shown). This observation is in agreement with leg and head tissues being mostly used for isolating genetic material. In testing for *Wolbachia* infection, we extracted genomic DNA from a limited number of available abdominal samples and performed sequencing of PCR amplicons using *Wolbachia*-specific primers (Fig. S3). *Wolbachia* amplicons suggestive of infection were detected in all samples tested, independently of mitochondrial haplotype, supporting no influence of *Wolbachia* infection on reproductive isolation of clustered populations. Further, lack of clear correspondence between genome-based and mitochondria-based haplotypes suggest on-going hybridization between C- and R-strains, consistent with detection of two potential hybrids among our 54 samples. Consistent lack of concordance between the nuclear and mitochondrial F_st_ values suggests that biological mitochondria-nuclear interactions are not maintained, further supporting that the samples analyzed are part of a panmictic population with little to no population structure.

Previous reports comparing genetic diversity in laboratory-reared insects to wild populations mostly reported loss of allele number and heterozygosity during adaptation to mass rearing [53, 54], although in some examples high genetic variability remained [55], even in long-established lines [56]. We compared diversity in two long-established (>20 years) *S. frugiperda* laboratory lines. Both laboratory lines, Benzon (USAMSs1) and SIMRU (USAMSs2), originated from field collections in Mississippi and they were the least differentiated from the TX/MN field collections. Interestingly, more recently laboratory populations from NC and FL [5] presented lower F_st_ values compared to field samples. This moderate differentiation between laboratory and field samples could be indicative of good culture management involving maintenance of large populations and/or regular introgression of field individuals. One possibility could also be that some of the field samples compared display low diversity, yet this is unlikely given samples from Central and South America (for Texas population) and the Caribbean (for Florida populations) based on current *S. frugiperda* migratory models [25]. However, these observations of moderate differentiation in laboratory and field populations are in contrast to previous research on locusts where genetic differentiation was decreased in laboratory colonies compared to field samples [57]. It is important to consider that despite the lack of overall differentiation between laboratory and field *S. frugiperda*, differentiation probably occurs during adaptation to mass rearing, resulting in selection of specific alleles that may not be representative of field populations. Thus, while our observations suggest laboratory strains contain genetic diversity similar to field populations, laboratory strains may have specific allele frequencies that may not be representative of field populations.

Overall, this study provides a diverse characterization of *S. frugiperda* whole and mitochondrial genome diversity. Findings from the study support lack of clear *S. frugiperda* population structure among the locations sampled, with only mitochondrial genomes indicating two different maternal lines mostly separating host strains. The genomic resources generated allow further exploration of gene flow and how it may affect management of *S. frugiperda* as an expanding global superpest using laboratory-reared and field-collected individuals.

## Acknowledgements

This project was funded by Agriculture and Food Research Initiative Foundational Program competitive grant 2018-67013-27820 and Hatch Multistate NC-246 from the US Department of Agriculture National Institute of Food and Agriculture. We would like to thank Rodney Nagoshi for review and helpful comments.

## Figure legends

**Table 1:** Pairwise F_st_ values across all populations surveyed. NA was used when the samples overlap in the defined populations.

