## supplemental for "Whole genome comparisons reveal panmixia among fall armyworm (*Spodoptera frugiperda*) from diverse locations"

### Materials and Methods

#### Detection of *Wolbachia*

Total genomic DNA was extracted from the abdomen of individual *Spodoptera frugiperda* moths using the Blood and Tissue DNEasy Kit (Qiagen) following the manufacturer's protocol. Primers to amplify the *Wolbachia* surface protein (*wsp*) marker (Zhou et al., 1998) were *wsp* 81F (5'- TGG TCC AAT AAG TGA TGA AGA AAC - 3') and *wsp* 691R (5'- AAA AAT TAA ACG CTA CTC CA - 3'). Reactions included 2x PCR Master Mix (Invitrogen™ Platinum™ SuperFi™), 1 µM of each primer, 10 ng of gDNA and nuclease-free water to 25 µl of final volume. Amplification conditions in an Eppendorf Mastercycler Ep Gradient Thermal Cycler included initial denaturation at 98°C for 30 sec, followed by 35 cycles of denaturation at 98° for 10 sec, annealing at 55.9°C for 10 sec and extension at 72°C for 30 sec, finalized with a final extension step at 72°C for 30 sec. A positive control consisting of *Wolbachia pipientis* gDNA provided by The Wolbachia project (Vanderbilt University, TN, USA) was included in all experiments. Amplicons were checked through 1% agarose gel electrophoresis, purified and sequenced at the Sequencing Core facility at the University of Tennessee (Knoxville).

### Supplementary Tables

**Supplementary Table 1** - List of *Spodoptera frugiperda* samples sequenced and used in this work. Shown is information on the location where collected (or facility for laboratory reference strains), Cry1F resistance phenotype (u for unknown, s for susceptible, r for resistant), collection (F for field-collected, L for lab-reared), host strain based on *Tpi*183 marker (corn, rice, hybrid or unknown), and the stage used for genomic DNA purification.

| Sample Name | Country | City/County/Facility | Municipality/ State originally collected | Cry1F | Collection | Host strain | Stage |
|---|---|---|---|---|---|---|---|
| ArgBAu1 | Argentina | Buenos Aires | Buenos Aires | u | F | Corn | Moth |
| ArgJs1 | Argentina | Estacion Experimental Agroindustrial Obispo Colombres (EEAOC, Tucuman) | Humahuaca (Jujuy) | s | L | Corn | Moth |
| ArgSFu1 | Argentina | San Justo | Santa Fe | u | F | Corn | Moth |
| ArgXXr1 | Argentina | Overo Pozo | Tucuman | r | L | Corn | Moth |
| ArgXXr2 | Argentina | La Cocha | Tucuman | r | L | Corn | Moth |
| BraBAr1 | Brazil | Luís Eduardo Magalhães | Bahia | r | L | Corn | Larva |
| BraBAr2 | Brazil | Luís Eduardo Magalhães | Bahia | r | L | Corn | Larva |
| BraBAr3 | Brazil | Luís Eduardo Magalhães | Bahia | r | L | Corn | Larva |
| BraBAr4 | Brazil | Luís Eduardo Magalhães | Bahia | r | L | Corn | Larva |
| BraBAr5 | Brazil | Luís Eduardo Magalhães | Bahia | r | L | Corn | Larva |
| BraMGr1 | Brazil | Viçosa | Minas Gerais | r | L | Corn | Larva |
| BraMGr2 | Brazil | Viçosa | Minas Gerais | r | L | Corn | Larva |
| BraMGr3 | Brazil | Viçosa | Minas Gerais | r | L | Corn | Larva |
| BraMGr4 | Brazil | Viçosa | Minas Gerais | r | L | Corn | Larva |
| BraMGr5 | Brazil | Viçosa | Minas Gerais | r | L | Corn | Larva |
| BraSPr1 | Brazil | Casa Branca | Sao Paulo | r | L | Corn | Larva |
| BraSPr2 | Brazil | Casa Branca | Sao Paulo | r | L | Unknown | Larva |
| BraSPr3 | Brazil | Casa Branca | Sao Paulo | r | L | Corn | Larva |

| | | | | | | | |
|---|---|---|---|---|---|---|---|
| KenXXu1 | Kenya | International Centre of Insect Physiology and Ecology (ICIPE, Nairobi) | Siaya and Homa Bay counties | u | L | Corn | Moth |
| KenXXu2 | Kenya | International Centre of Insect Physiology and Ecology (ICIPE, Nairobi) | Siaya and Homa Bay counties | u | L | Corn | Moth |
| KenXXu3 | Kenya | International Centre of Insect Physiology and Ecology (ICIPE, Nairobi) | Siaya and Homa Bay counties | u | L | Corn | Moth |
| PueGUr1 | Puerto Rico | | Guayama | r | F | Corn | Larva |
| PueLAu1 | Puerto Rico | | Lajas | u | F | Corn | Moth |
| PueSIu1 | Puerto Rico | | Santa Isabel | u | F | Rice | Moth |
| USAFLr1 | USA | Collier County | Florida | r | L | Corn | Larva |
| USAFLr2 | USA | Collier County | Florida | r | L | Corn | Larva |
| USAFLu1 | USA | Belle Glade (Palm Beach County) | Florida | u | F | Corn | Moth |
| USAFLu10 | USA | Hague (Alachua County) | Florida | u | F | Corn | Moth |
| USAFLu11 | USA | Hague (Alachua County) | Florida | u | F | Corn | Moth |
| USAFLu12 | USA | Hague (Alachua County) | Florida | u | F | Corn | Moth |
| USAFLu2 | USA | Belle Glade (Palm Beach County) | Florida | u | F | Rice | Moth |
| USAFLu3 | USA | Miami (Miami-Dade County) | Florida | u | F | Corn | Moth |
| USAFLu4 | USA | Miami (Miami-Dade County) | Florida | u | F | Corn | Moth |
| USAFLu5 | USA | Miami (Miami-Dade County) | Florida | u | F | Hybrid | Moth |
| USAFLu6 | USA | Miami (Miami-Dade County) | Florida | u | F | Corn | Moth |
| USAFLu7 | USA | Miami (Miami-Dade County) | Florida | u | F | Corn | Moth |
| USAFLu8 | USA | Hague (Alachua County) | Florida | u | F | Corn | Moth |
| USAFLu9 | USA | Hague (Alachua County) | Florida | u | F | Corn | Moth |

| | | | | | | | |
|---|---|---|---|---|---|---|---|
| USAMDu1 | USA | Jarretsville (Harford County) | Maryland | u | F | Rice | Moth |
| USAMDu2 | USA | Jarretsville (Harford County) | Maryland | u | F | Rice | Moth |
| USAMNu1 | USA | Rosemount (Dakota County) | Minnesota | u | F | Corn | Moth |
| USAMNu2 | USA | Rosemount (Dakota County) | Minnesota | u | F | Corn | Moth |
| USAMSs1 | USA | Benzon Research Inc (Carlisle, PA) | Mississippi | s | L | Corn | Moth |
| USAMSs2 | USA | USDA-ARS Southern Insect Management Reseacrh Unit (SIMRU, Stoneville) | Mississippi | s | L | Corn | Moth |
| USANCr1 | USA | Hyde County | North Carolina | r | L | Corn | Moth |
| USASCu1 | USA | Charleston (Charleston County) | South Carolina | u | F | Corn | Moth |
| USASCu2 | USA | Charleston (Charleston County) | South Carolina | u | F | Corn | Moth |
| USATNu1 | USA | Crossville (Cumberland County) | Tennessee | u | F | Rice | Moth |
| USATNu2 | USA | Crossville (Cumberland County) | Tennessee | u | F | Rice | Moth |
| USATXu1 | USA | Corpus Christi (Nueces County) | Texas | u | F | Unknown | Moth |
| USATXu2 | USA | College Station (Brazos County) | Texas | u | F | Rice | Moth |
| USATXu3 | USA | College Station (Brazos County) | Texas | u | F | Rice | Moth |
| USATXu4 | USA | Corpus Christi (Nueces County) | Texas | u | F | Hybrid | Moth |
| USATXu5 | USA | Lubbock (Lubbock County) | Texas | u | F | Hybrid | Moth |
| USATXu6 | USA | Lubbock (Lubbock County) | Texas | u | F | Corn | Moth |

**Supplementary Table 2 -** Average Mash distances based on country of origin based on all 55 samples

| Country of origin | Argentina | Brazil | Kenya | Puerto Rico | USA |
|---|---|---|---|---|---|
| **Argentina** | | 0.043 | 0.044 | 0.044 | 0.044 |
| **Brazil** | 0.043 | | 0.043 | 0.045 | 0.045 |
| **Kenya** | 0.044 | 0.043 | | 0.043 | 0.043 |
| **Puerto Rico** | 0.044 | 0.045 | 0.043 | | 0.045 |
| **USA** | 0.044 | 0.045 | 0.043 | 0.045 | |

### Supplementary figures

**Figure S1**: Geographical distribution based on all the 55 *S. frugiperda* samples used in the study with dots proportional to the number of samples collected for each location.

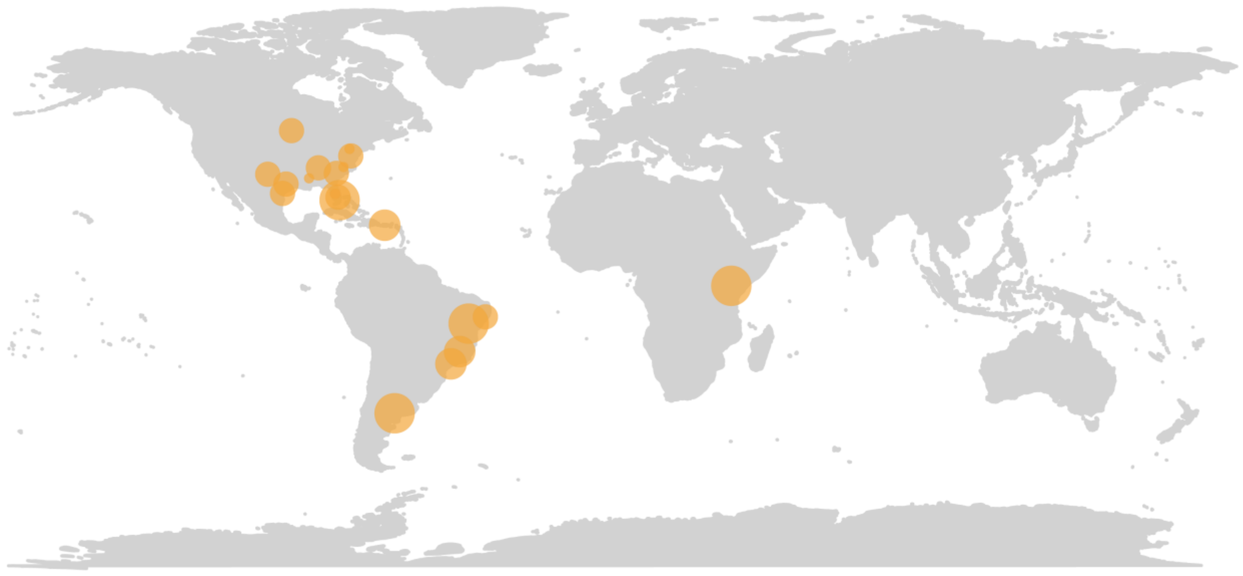

**Figure S2**: Complete clustering on Mash distances heatmap of Mash distances across 55 samples colored by host strain and rooted with *Spodoptera litura*.

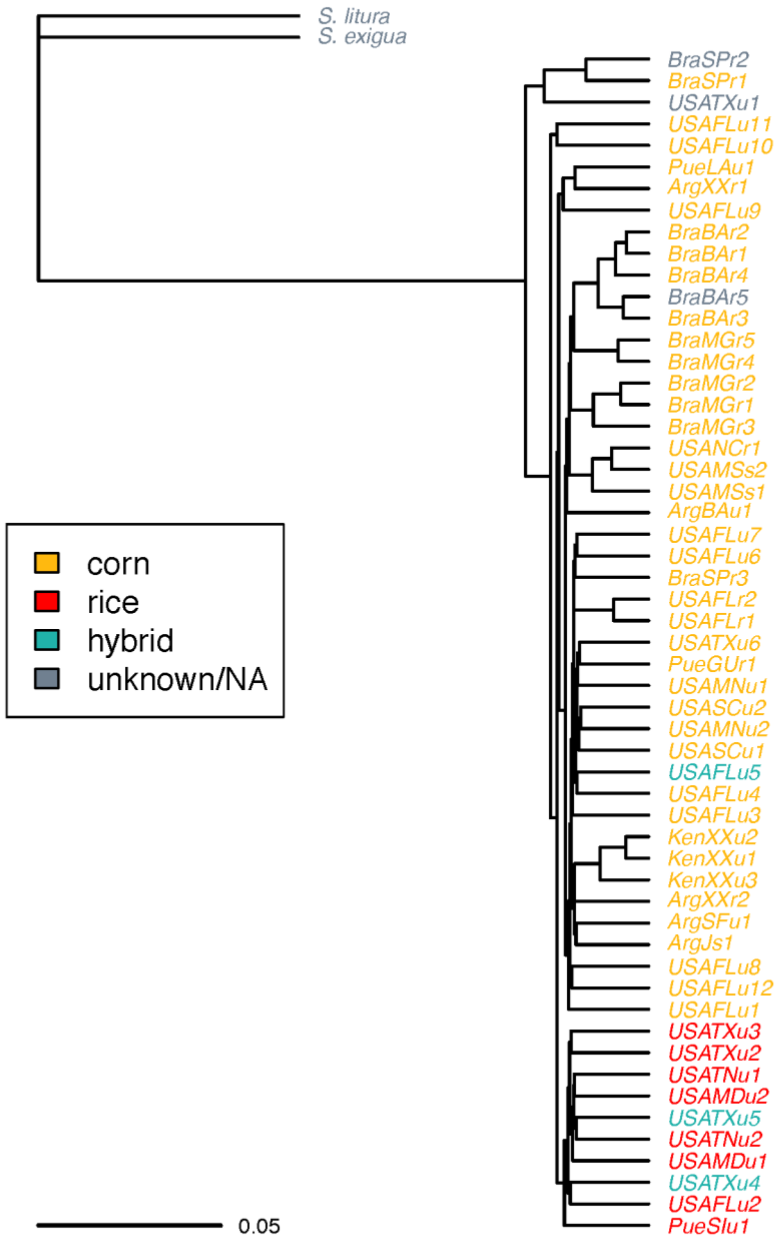

**Figure S3**: Amplification of a fragment of the *wsp* gene (610 bp) from *S. frugiperda* moths. A positive control (*Wolbachia pipiens*) was also included (+C). Samples shown ArgXXr2 (lane 1), PueSIu1 (lane 2), USAFLu8 (lane 3), USAFLu10 (lane 4), USAFLu12 (lane 5), USAMDu2 (lane 6), USATXu5 (lane 7) and *W. pipiens* (lane 8). Sequenced amplicons matched with 99.8% identity to *Wolbachia* endosymbiont of *Nasonia vitripennis* seqvar1 Nvit outer surface protein (*wsp*) gene (GenBank accession number DQ380865).

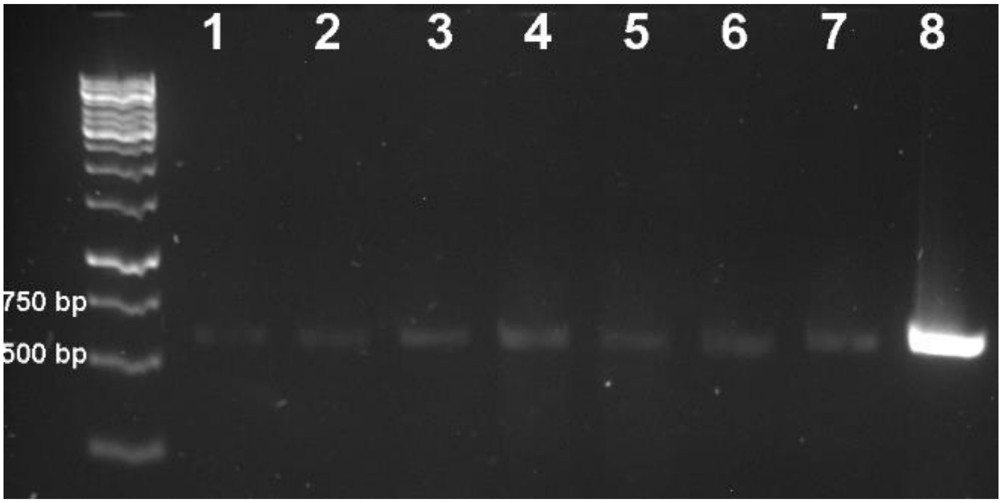
